# In harmony? A scoping review of methods to combine multiple 16S amplicon data sets

**DOI:** 10.1101/2025.02.11.637740

**Authors:** Zone Li, Yingan Chen, Yage Sun, Kristen McArthur, Megan Carnes, Tiange Liu, Noel T. Mueller, Grier P. Page, Lori Rossman, Ekaterina Smirnova, Julie D. White, Amii M. Kress, Justine W Debelius

**Affiliations:** Department of Epidemiology, Johns Hopkins Bloomberg School of Public Health; Baltimore, MD, USA; Department of Epidemiology, Colorado School of Public Health, Lifecourse Epidemiology of Adiposity and Diabetes (LEAD) Center, University of Colorado, Anschutz Medical Campus, Aurora, CO, USA; Cutaneous Translational Research Program, Department of Dermatology, Johns Hopkins School of Medicine, Baltimore, MD, USA; GenOmics and Translational Research Center, RTI International; Research Atlanta, GA, USA; Division of Women’s Health, Department of Medicine, Brigham and Women’s Hospital and Harvard Medical School; Boston, MA, USA; Department of Pediatrics, Section of Nutrition, University of Colorado, Anschutz Medical Campus, Aurora, CO, USA; Welch Medical Library, Johns Hopkins School of Medicine, Baltimore, MD; Department of Biostatistics, Virginia Commonwealth University, Richmond, VA

**Keywords:** meta research, microbiome, meta-analysis, scoping review, 16S sequencing, human microbiome

## Abstract

Robust evidence on relationships between the human microbiome and health are critical for understanding and improving the human condition. However, there is little information about methodological approaches to combine and analyze multiple microbiome data sets. To address this gap, we conducted a scoping review of studies that combine sequencing data from multiple data sets to understand the study objectives, data sources and selection, feature -table assembly methods, and analyses. References were identified through a systematic search of literature published between 2011 and 2022. Our final review included 60 articles. Despite the wide-spread use of the word “meta-analysis,” we found that only 24 studies used a systematic process to select their data sets, suggesting multiple meanings of the term within the field. While more than two thirds of studies used at least one publicly available data source, 19 had to request data from the original authors. Most studies (60%) combined data sets from multiple disjoint hypervariable regions. The number of hypervariable regions combined was not associated with the table construction method, but feature table construction method and the number of hypervariable regions influenced analytical resolution. Our results suggest the microbiome community needs to examine the use of terminology and analytic approaches for combining data sets; that additional work is needed to explore the impact of data source on bias in combined studies; and invite an independent evaluation of the methods used for feature table construction across disjoint regions.

## Introduction

The human microbiome is hypothesized to play a critical role in health, affecting development, metabolism, the immune system, and cognition (1–3). Many of these conclusions are based on small, single center studies, limiting their generalizability. Understanding the role of the microbiome in health and moving forward interventions requires consistent findings. However, evidence synthesis studies, such as systematic reviews and meta-analyses based on summary statistics, often fail to find evidence supporting the association of common taxa with the outcome of interest, or they find common taxonomic associations among such broad clades (i.e. class, phylum) that mechanistic inference is limited (4–7). Few synthesis studies mention the potential for the methodological choices in the data sets combined (e.g. differences in extraction, taxonomic annotation, or statistical testing) to influence the observed results, affecting the ability to make an overall generalization about the relationship between the microbiome and a disease of interest (4–6). For example, there is limited interoperability between taxonomic databases used for 16S rRNA annotation, meaning the same sequence may be assigned to a different clade if a different database is used (8–11).

The rapid changes in organism naming conventions and taxonomy-phylogeny relationships in databases further exacerbate these issues (11–15). Furthermore, recent work has suggested common tools to test for differentially abundant taxa between groups produce disturbingly different results on the same data set (16). The failure to consider or discuss technical differences casts doubt on the conclusions of systematic reviews and statistical meta-analyses, as the included studies are often not directly comparable. In other words, it is possible that inconsistencies noted in the systematic reviews may be due to differences in bioinformatic choices across studies, rather than true biological variation between populations. (8–11).

The best solution to evidence synthesis may therefore be a combined re-analysis of previously published primary data (sequences and participant data) using a common bioinformatic pipeline and statistical approach. This proposition is complicated by heterogeneity in extant data sets and published studies. Each step in sample collection, processing, amplification, sequencing, and bioinformatics biases the observed microbial community (16,17). The selected method often reflects a set of value propositions that may be appropriate for the analysis being performed but might not be applicable under other conditions. For example, microbiome characterization using the 16S rRNA gene often relies on short amplicons that target a fragment of the full gene (18,19). Primer pairs used to amplify different regions have different biases, specificities, and taxonomic resolutions; the “optimal” primer pair for a study depends on the environment being studied (18–21). Furthermore, there is a consensus against standardization as this will enshrine a single set of biases, rather than allowing more robust triangulation of results (17).. Thus, re-analysis of primary data necessitates harmonization across multiple non-overlapping fragments of the 16S rRNA gene. There are multiple possible bioinformatic strategies that might be used to address the challenge of harmonization, each with benefits and drawbacks. We highlight five potential approaches in Table 1, based on the feature table assembly techniques used in microbiome research. While these approaches are common in microbiome analysis, little work has been done to characterize how researchers are applying these approaches in the context of combining multiple data sets.

**Table 1.** Approaches to 16S rRNA gene harmonization.

|  | **Description and taxonomy** | **Feature identification** | **Beta (between sample)**<br>**Diversity based comparisons** | **Feature-based**<br>**comparisons (Differential**<br>**abundance, machine**<br>**learning)** |
| --- | --- | --- | --- | --- |
| Amplicon<br>Sequence<br>Variants<br>(ASVs) | Sequencing and PCR noise are removed or corrected from the sequence, resulting in a single nucleotide resolution (71).<br>Taxonomy is assigned to the representative sequence using a variety of algorithms including naïve Bayesian classification, BLAST, or phylogenetic placement (72–75). | Features are represented by a sequence, dependent on the position on the 16S gene and trimming length (71) .<br>Representative ASV sequences are not comparable across regions. | Data collapsed to a common taxonomic description can be compared across regions (i.e. genus level) using metrics that do not require a phylogenetic tree.<br>Janssen et al (41) proposed phylogenetic scaffolding could combine ASVs with different hypervariable regions for phylogenetic metrics. | Data collapsed to a common taxonomic description can be compared across regions (e.g. genus level). |
| Closed<br>reference<br>clustering<br>into<br>Operational<br>taxonomic<br>units (OTUs) | Sequences are clustered against a reference database at some level of comparison to the reference sequence. Reads which do not match the reference are discarded (76)<br>Taxonomy is inherited from the reference database (76). | Features are identified based on the reference sequence, which has a consistent identifier across the regions in the database where it is present (76,77) | The reference OTU ID functions as a common identifier, and data can be compared directly at the feature level using phylogenetic or nonphylogenetic metrics (77).<br>Data can also be collapsed to a common taxonomic description (i.e. genus level) with non-phylogenetic metrics if necessary | The reference OTU ID functions as a common identifier, and data can be compared directly at the feature level (77).<br>Data can also be collapsed to a common taxonomic description (i.e. genus level) for comparison |

|  | **Description and taxonomy** | **Feature identification** | **Beta (between sample)**<br>**Diversity based comparisons** | **Feature-based comparisons (Differential abundance, machine learning)** |
| --- | --- | --- | --- | --- |
| de novo OTU clustering | Sequences are clustered based on similarity threshold. All high-quality reads are retained (76).<br>Taxonomy is assigned via a naïve Bayesian classifier, BLAST-ing sequences against a reference, or other methods (72,75) | Features are identified based on a centroid sequence from the cluster.<br>Centroid sequences from the de novo clustering are not comparable between regions | A de novo phylogenetic tree cannot be compared across hypervariable regions so phylogenetic metrics are not comparable.<br>Data can be collapsed to a common taxonomic description (i.e. genus level) with non-phylogenetic metrics. | Data can also be collapsed to a common taxonomic description (i.e. genus level) for comparison |
| Open reference OTU clustering | Sequences are clustered against a reference database, and discarded sequences are then clustered against themselves (76,78)<br>Taxonomy is inherited from the reference database for reference-assigned sequences and using a naïve Bayesian classifier for the de-novo clusters (76,78). | Features are identified based on the reference database or the centroid sequence.<br>The reference-based clusters are comparable across regions, the de novo centroids are not, making it difficult to compare the entire data set across regions. | Data can be collapsed to a common taxonomic description (i.e. genus level) with non-phylogenetic metrics. | Data can be collapsed to a common taxonomic description (i.e. genus level) with non-phylogenetic metrics. |
| MiAmpLiPi (70) | Sequences are denoised, placed into a reference phylogenetic tree, and then binned into phylotypes based on phylogenetic similarity thresholds<br>Taxonomy is inherited from the full-length reference sequences used to generate the tree. | Features are identified based on a common phylotype ID. | Phylotype IDs are directly comparable across hypervariable regions and can be used as features for phylogenetic and non-phylogenetic metrics. | Phylotype IDs are directly comparable across hypervariable regions and can be used as features for differential abundance or machine learning. |

To address this gap, we conducted a scoping review of studies that combined primary 16S rRNA data sets and analyzed them together. Our goals were to describe the studies and how they selected and acquired specific data sets, how 16S data were harmonized across data sets, and how the harmonization approach influenced statistical decisions.

## Methods

### Search Strategy and Screening

We did not pre-register a study protocol.

We were interested in three concepts in our search: the microbiome; data combination; and sequencing of the 16S rRNA gene. Full search terms used for identifying publications are described in Supplemental File 1. References were limited to those published between 2011 and June 2022. We considered Andersson et al, 2008 (22) to be the first human microbiome study with next generation sequencing techniques. Our 2011 start year therefore allowed time for studies combining multiple 16S sequencing data sets to be published. References were identified through a systematic search of PubMed, Embase, Scopus, Web of Science, and the Cochrane library, performed on June 9, 2022.

References were screened through the Covidence webtool (Melbourne, Australia) with a two-step process. References were excluded at the title and abstract stage if they were obviously a commentary, perspective, or erratum; did not include primary sequence analysis (e.g. systematic reviews); or represented a single 16S sequencing data set. Title-abstract screening was performed independently by two reviewers (YC and TL, MC, KM, or JWD). In full text screening, eligible articles were peer reviewed studies, written in English, focused on the combination of 16S rRNA sequencing data of microbial communities from multiple sources (labs or published studies). Full text screening was performed independently by two reviewers (YC and MC, KM, JDW, or JWD). In both title-abstract and full-text screening, conflicts were resolved by a senior third-party reviewer (JWD, JDW, or MC) and through group discussion and mutual consensus. During full text screening, studies were categorized by primary environment (human, animal, built environment, wastewater, ocean, soil, plant, or other) and only studies including human samples were included for data extraction.

### Data Extraction

From each article, we extracted information on the goal of combining the 16S data, characteristics of the data sets included, how feature tables were constructed, and how statistical analyses were conducted. Information was extracted using a structured form in a Qualtrics web survey (Qualtrics, Provo, Utah, USA). The full extraction form can be found on Zenodo (doi: 10.5281/zenodo.14767165) (23).

Article extraction was performed by two of four independent reviewers (ZL, YC, YG, or JWD). Reviewers trained on a common set of four articles and discrepancies were discussed. Inter-reviewer correlations were checked regularly, and all four reviewers met to discuss questions about the papers. Conflicts were resolved by a third reviewer who had access to the responses of the previous two reviewers. The results of the data abstraction were assembled using the pandas library (v1.2.5) in python 3.8.11 (24,25).

### Synthesis of Results

All analyses were conducted in R (v4.2.2) using the tidyverse data ecosystem (v2.0.0) (26,27). Upset plots were constructed using the ComplexUpset library (v1.3.3); other figures were made using ggplot2 (v3.5.1) (28–30). Analyses were primarily descriptive. Associations between the number of hypervariable regions assessed, feature table construction method, and taxonomic level for analysis were tested with Fisher’s exact tests. An alpha value of < 0.05 was considered statistically significant.

### Code and Data Availability

The resolved, extracted data, a data dictionary, and the extraction form are available on Zenodo (10.5281/zenodo.14767165) (23). Analysis code used in this project is on GitHub (https://github.com/jwdebelius/MicroScoping) and Zenodo (10.5281/zenodo.14814589) (31).

## Results

Our review focused on articles which combined primary data form multiple 16S rRNA sequencing data sets. Using a systematic search strategy, we identified 946 references for evaluation, of which 696 were excluded at the title and abstract stage and an additional 139 were excluded at full text screening. Of the 111 articles that met the broader inclusion criteria, the 60 that included human microbiome data are presented in this paper (Figure 1, Table 2). The agreement between each pair of independent extractors was greater than 82.5%.

**Figure 1.**
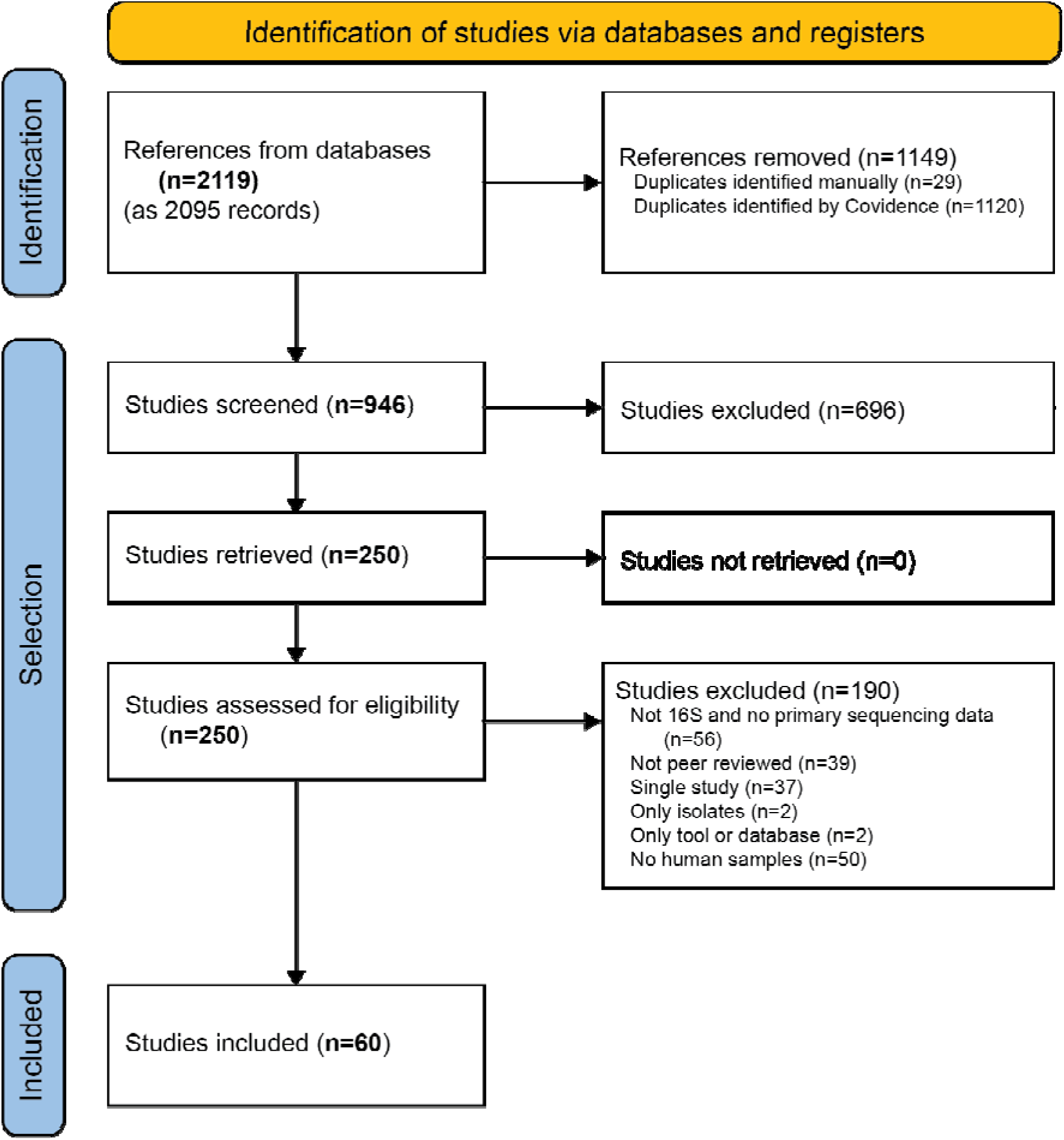
**Selection of studies included in the scoping review**

**Table 2.** Characteristics of studies included in this scoping review.

|  | Total N (%) |
| --- | --- |
| **Number of studies included** | 60 |
| **Analytical purpose of the combined analysis<sup>1</sup>** |  |
| Population synthesis | 53 (88) |
| Describing a core microbiome | 5 (8) |
| Replication and validation of previous findings | 5 (8) |
| Pan environmental comparisons, including for method validation | 3 (5) |
| **Scientific goal of the combined analysis<sup>1</sup>** |  |
| Describe the relationship between the microbiome and an outcome | 41 (68) |
| Describe the relationship between the microbiome and an exposure | 12 (20) |
| Studying community Assembly or Dynamics | 5 (8) |
| To characterize a unique microbiome | 6 (10) |
| Other<sup>2</sup> | 6 (10) |
| **Environment** |  |
| Human only | 54 (90) |
| Human and other animals | 2 (3) |
| Pan environmental surveys | 4 (7) |
| **Human body site(s) analyzed<sup>1</sup>** |  |
| Gut | 50 (83) |
| Oral | 9 (15) |
| Airway | 8 (13) |
| Urogenital | 7 (12) |
| Skin | 5 (8) |
| Pan environmental or includes non-animal hosts | 4 (7) |
| **Number of data sets included** |  |
| ≤ 5 | 22 (37) |
| 6-10 | 9 (15) |
| 11-15 | 13 (22) |
| > 15 | 13 (22) |
| Not reported | 3 (5) |
| **Number of samples included** |  |
| < 500 | 13 (22) |
| 500-1500 | 12 (20) |
| 1501-2500 | 10 (17) |
| >2500 | 14 (23) |
| Not Reported | 11 (18) |
| **Included secondary data set(s) for validation** | 8 (13) |
| **Clear systematic approach for data set gathering** | 25 (41.7) |

<sup>1</sup>Multiple responses possible; number of papers with this response
<sup>2</sup>Other purposes included pan environmental comparisons; methodological assessments; similarities between humans and mouse strains; and microbiome-metabolite associations

The included studies (n = 60) were published between 2011 and 2022, although more than three quarters of the articles were published after 2017 (23). Articles were published in a wide variety of journals, including microbiome-focused publications, journals without a field specific focus, and outcome focused journals. The studies were designed with several scientific goals in mind, although most studies focused on a microbiome-outcome (n=41, 68%) or microbiome-exposure (n=12, 20%) association (Table 2). The analytical purpose of most studies was to combine multiple data sets to describe a single population (n=53 85%); four of these (7% of total) also tried to define a core microbiome, the common set of features for a particular environment. Five articles combined data sets with the intention to validate findings, such as the results of a classifier (8%) (32–36). Ninety percent (n=54) of studies focused on human samples only; two included samples from non-human animals (e.g. mice) and four were pan environmental comparisons which included human samples. Forty-two (70%) studies exclusively focused on the gut microbiome; three (5%) studies considered the gut in combination with one body site. Nine studies (15%) focused on other body sites, such as the oral, airway, urogenital, or skin microbiome, either alone or in combination. Five studies (8%) included samples from 3 or more body sites. The studies included between one and 2,462 data sets with a median of 10 (Interquartile range [IQR] 4 - 15) primary data sets; three papers did not explicitly report the number of data sets included. While 11 (18%) of studies did not report the number of samples used, most studies that did report samples sizes included ≥ 500 samples. Eight studies included validation data sets, with a median of 1.5 data sets (range 1 - 5). The two studies that included only one data set in their primary analysis also included at least one validation data set.

### Data set Selection and Description

We also abstracted information about how the studies identified data sets to include in their analysis, and how they described these data sets in the text. For this section, we focused on the 51 papers that combined data sets to describe a population or to test a hypothesis about the microbiome and specific exposures or health outcomes. Of the 51 relevant articles, 46 had the words “meta analyses” or “meta-analysis” in either the title or abstract. Less than half (n= 24; 47%) reported a systematic approach for data set identification and inclusion. Of these, 20 detailed their search strategies, 13 incorporated a PRISMA diagram for data set selection, and all 24 specified their inclusion criteria. Of the 51 articles, 39 (77%) included basic descriptive information on the source population. However, technical parameters describing the microbiome sample collection approach and extraction protocols used by combined data sets were rarely presented, with 10 (20%) and 11 (22%) articles providing sample collection and extraction details, respectively. In contrast, hypervariable region(s) were described for 43 (84%) and sequencing platform(s) were described in 37 (73%) of the 51 studies.

### Data set Sources

The microbiome community has emphasized the importance of publicly available data to support robust and reproducible research (37,38). Therefore, we were interested in where studies found their data sets (Figure 2). The majority (n=43; 72%) used at least one publicly available data source, such as the European Nucleotide Archive (ENA), Sequence Read Archive (SRA), Metagenomics – Rapid Annotation using Subsystem Technology (MG-RAST) server, and QIIME-DB, now Qiita. We note that sequences are mirrored between ENA and SRA (39). There were 33 (55%) studies that used closed data sources, including data sets from the authors of the combined study or requests to the authors of the original data set. We found 13 (21.7%) studies with other data sources such as the China National GeneBank DataBase (CNGB), GitHub, the American Gut Project, Dryad, ImmPort Shared Data, Mendeley Data, Chaffron data set, DNA Data Bank of Japan (DDBJ), Database of Genotypes and Phenotypes (dbGaP), and the Microbiome CRC Biomarker Study. Eight (13.3%) articles had one or more data set in their studies coming from sources that were not described or unclear. Overall, we found 17 (28%) studies that used only open data sources, while 19 (32%) studies required the study authors to request additional information about a previously published data set.

**Figure 2.**
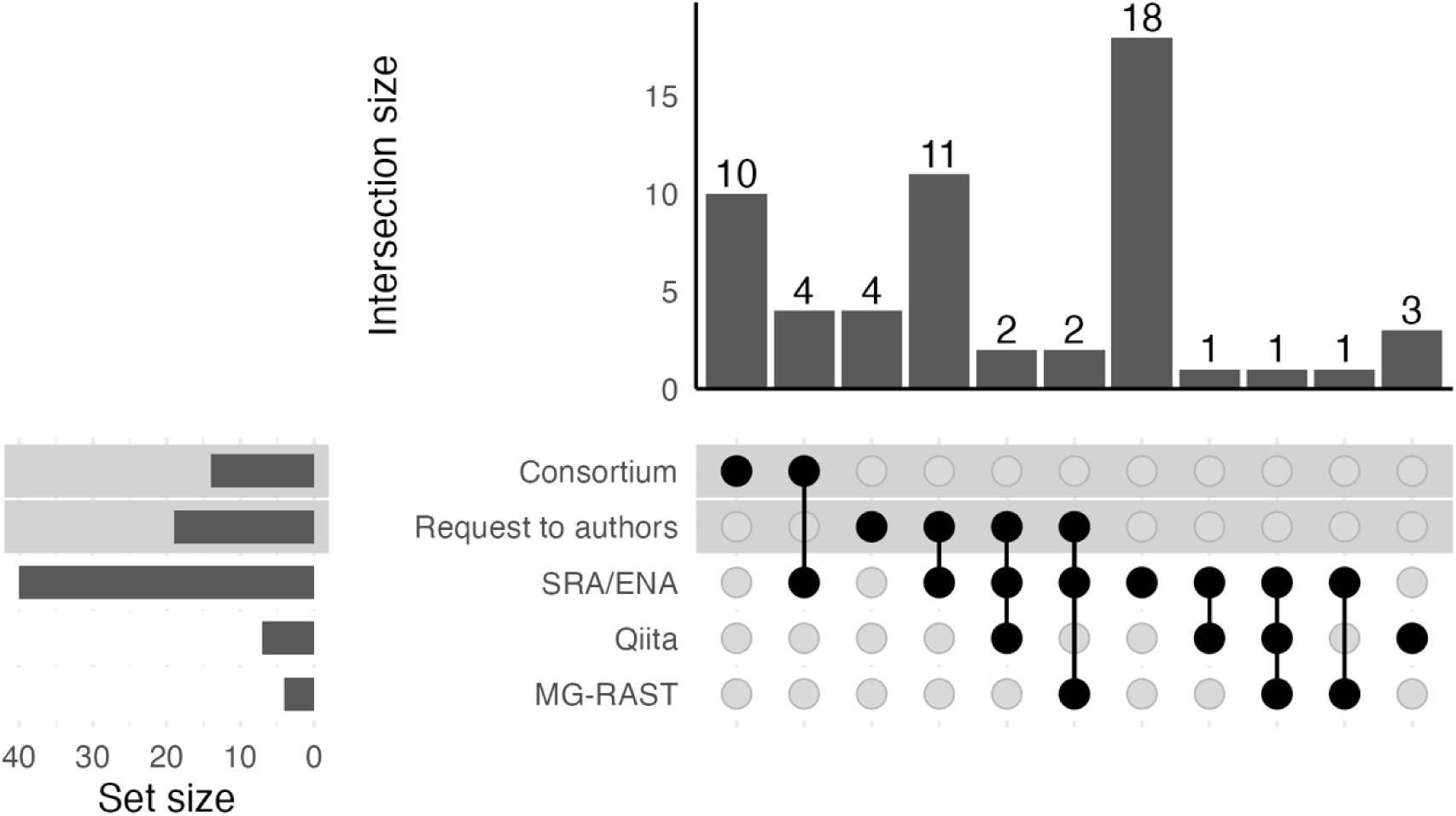
Upset plot comparing the number of data set sources used by study authors. The horizontal bar plot at the left gives the total number of studies which included a data set from a given source; the vertical bar plot shows the number of studies with for each intersection of data set sources. Closed data sources have a shaded background.

### Sequencing Platforms and Hypervariable Regions

The technical parameters used to sequence the data in our included studies were heterogeneous, which likely influences how 16S data were harmonized to produce a common feature table. While factors like sample collection and extraction are known to create biases in observed microbial communities (19), they should not substantially influence bioinformatic processing. We anticipated sequencing platform and hypervariable regions would influence data set processing and data harmonization (18). Among the 60 studies reviewed, 14 (23%) did not specify their sequencing platforms. A total of 28 (47%) included data sets amplified on the Roche 454 pyrosequencer; 40 (67%) included data sets amplified with Illumina and five studies (8%) had data sets amplified with Ion Torrent. Table 3 summarizes the observed combinations of sequencing platforms.

**Table 3.** Combination of Sequencing Platforms analyzed in different studies.

| **Combination of Sequencing platforms** | **Total (%)**<br>**(n = 60)** |
| --- | --- |
| Illumina only | 15 (25%) |
| 454 pyrosequencing only | 4 (7%) |
| Illumina and 454 sequencing | 21 (35%) |
| Illumina and Ion Torrent | 2 (3%) |
| Illumina, 454 and Ion Torrent | 2 (3%) |
| 454 and Ion torrent | 1 (2%) |
| Missing | 14 (23%) |

Previous research has shown that amplicons from different hypervariable regions within the same organism are not directly comparable, due to both the underlying biological functions of the 16S rRNA gene and any technical biases introduced during DNA amplification, sequencing, and bioinformatic processing (18). Additionally, the hypervariable region can have a large impact on the observed community, often exceeding the effect of biologically relevant factors (40,41). Over 80% (n=50) of the included studies reported the hypervariable regions amplified in their data sets (Figure 3). We found 14 (23%) studies included data from only one hypervariable region; 14 (23%) used data sets which targeted two or three hypervariable regions, and 22 (37%) included data sets from four or more regions. The most common hypervariable region included was V4 which was included in 40 of the studies (80% of the 50 studies with described hypervariable regions). Primers targeting the V34 and V35 regions were also common, present in 29 and 17 studies, respectively.

**Figure 3.**
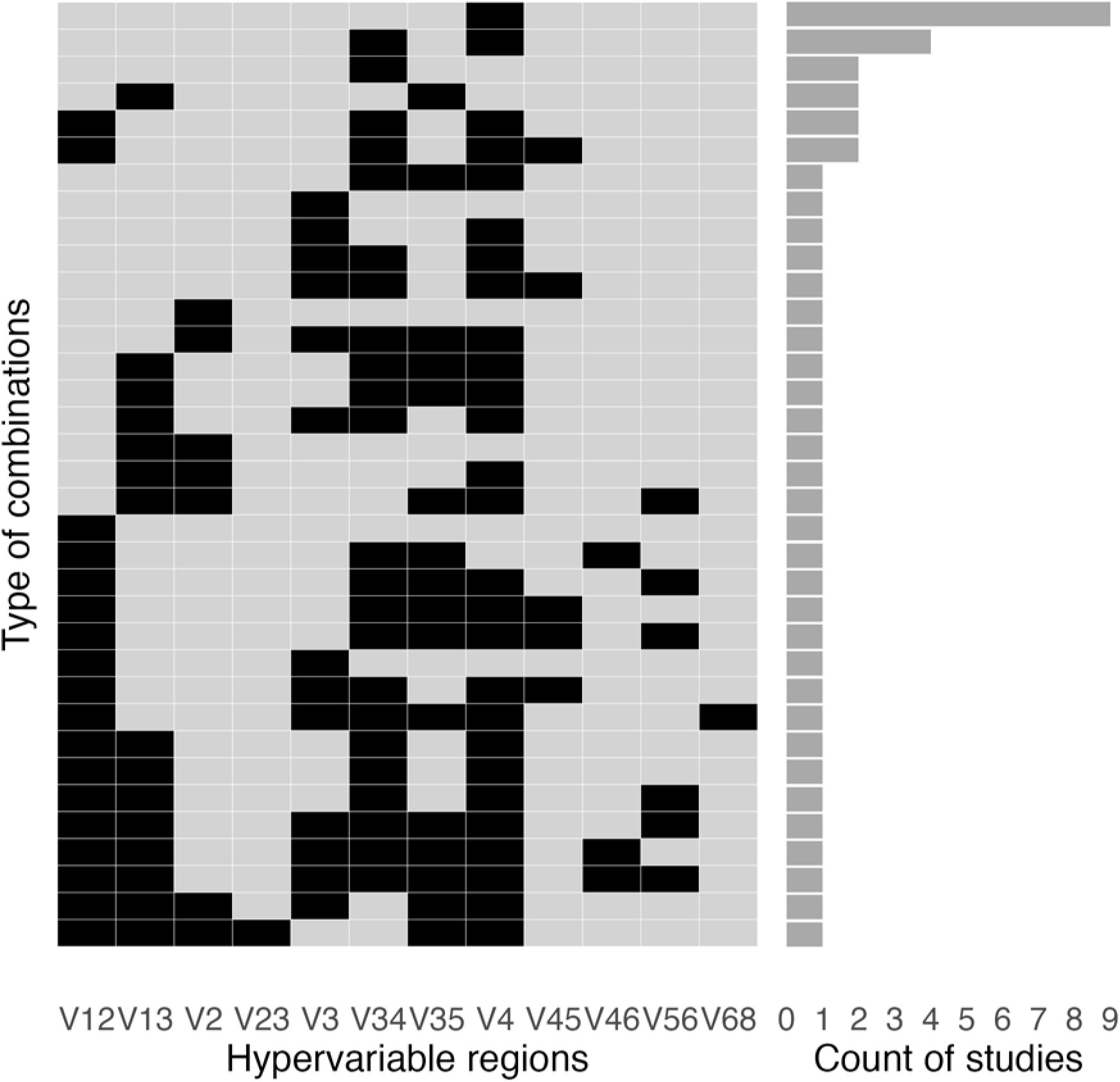
Hypervariable region combinations included in different studies. The matrix at the left shows the regions amplified by data sets included in each study; the bar chart at the right describes the number of studies which included that combination of hypervariable regions.

### Feature Table Construction Methods

Combining 16S sequencing data across multiple, disjoint hypervariable regions is challenging. One major goal in feature table construction is to harmonize the regional sequence identifiers in such a way that the same organism shares a common identifier across disjoint regions. This decision involves a balance between retaining high sequence-based resolution, which provides more precision and allows the use of phylogenetic techniques (42); or working with data collapsed to a common taxonomic annotation, a choice which may decrease data set specific technical biases but may also obfuscate true ecological differences between microbial communities (43) (Table 1). We were unable to identify the feature table construction method for seven studies, either because they did not report one or because they did not use consistent method for table construction across data sets. Of the remaining 53 studies, a total of 12 (20%) studies analyzed amplicon sequence variants (ASVs), 18 (30%) used closed reference operational taxonomic units (OTU) clustering, eight (13%) studies clustered their data into *de novo* OTUs, and one (2%) used open reference clustering. Ten (17%) studies used some type of OTU clustering, but the specific method was not clearly described. Other approaches included the use of a proprietary pipeline (44) and direct taxonomic classification without denoising or clustering (45). In a replication study, open reference clustering was used on the primary data set and the replication data sets performed closed reference clustering against the open reference centroids (35). Our results therefore suggest that while closed reference clustering was the most common method, other approaches were applied.

### Analyses conducted

We additionally characterized the types of analyses performed by each study. Analyses were clearly described in 55 of 60 studies (Figure 4). Eleven (20%) included descriptive analyses like taxa plots; 44 (80%) considered alpha (within sample) diversity; 41 (74%) included beta or between sample diversity; 44 (80%) performed some kind of differential abundance testing to look for individual organisms that differed between groups; and 21 (38%) used machine learning. Additionally, eight (14%) studies looked for a “core” microbiome, or a common set of features shared across an environment or disease, and four (7.3%) performed co-occurrence networking. Two studies constructed community state types (46,47). Studies performed a median of three types of analyses (IQR 2 - 4 analyses) with 38 studies using three or more analytical techniques. The combination of alpha diversity, beta diversity, and differential abundance was most common (n=17), with an additional 14 studies performing these three analyses in addition to machine learning to predict an outcome or exposure.

**Figure 4.**
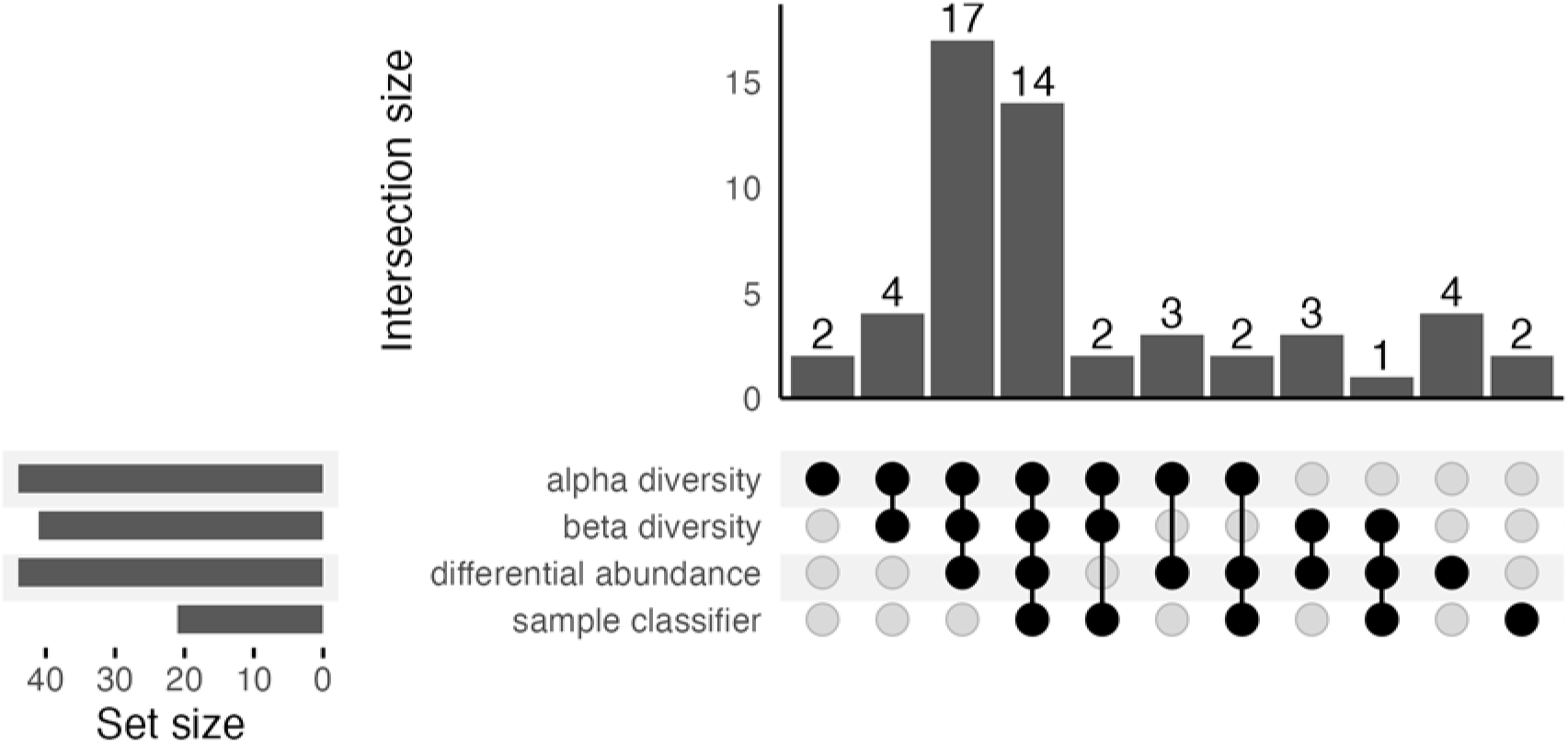
Analyses conducted by individual studies. Upset plot shows the number of studies which conducted each analysis, and the number of studies which conducted each combination. Five studies which did not report their analyses conducted are excluded.

Among the four most common analyses (alpha diversity, beta diversity, differential abundance, and machine learning), we also assessed the lowest taxonomic level where the analysis was performed, from feature (ASV or OTU) to phylum (Table 4). Overall, we found that alpha and beta diversity were commonly conducted at the feature level, differential abundance was almost evenly split between the feature level and the genus level.

**Table 4.** Lowest taxonomic level used for analysis by study and analysis type.

| **Lowest taxonomic level** | **Alpha Diversity (n=44)** | **Beta Diversity (n=40)** | **Differential Abundance (n=44)** | **Sample Classification (n=21)** |
| --- | --- | --- | --- | --- |
| OTU or ASV | 29 (66%) | 30 (75.0%) | 19 (43%) | 12 (57%) |
| Species | 5 (11%) | 3 (7%) | 5 (11%) | 0 ( 0%) |
| Genus | 4 (9%) | 4 (10%) | 18 (41%) | 7 (33%) |
| Family | 0 (0.0%) | 1 ( 3%) | 0 (0%) | 0 ( 0%) |
| Phylum | 0 (0.0%) | 0 ( 0%) | 2 (5%) | 0 ( 0%) |
| Unclear | 6 (14%) | 2 ( 5%) | 0 (0%) | 2 (10%) |

### Table Construction Method and Taxonomic Resolution

This scoping review also aimed to evaluate the relationship between the number of hypervariable regions used, feature table construction method, and taxonomic collapse. Researchers could solve combining common features across regions multiple ways (Table 1). Working with one hypervariable region (or one primer pair) facilitates easy harmonization at the feature (ASV or OTU level) using any feature table construction method. To combine multiple hypervariable regions, some kind of scaffolding is necessary. This can either come as closed reference OTU clustering, where the reference identifier becomes the scaffold, or by collapsing the data to a specific taxonomic level and comparing, for example, the genus level abundances. We hypothesized that studies which used feature table construction methods that do not allow a direct comparison between features from different hypervariable regions (e.g., de novo OTU clustering; denoising to ASVs) would be more likely to perform analyses that allowed feature-based comparisons after collapsing the data (e.g., differential abundance; Table 1). We restricted our analysis to the 32 studies which (1) used either closed reference clustering, de novo OTU clustering, or denoised to ASVs, (2) provided information about the number of hypervariable regions, and (3) analyzed alpha diversity, beta diversity, or differential abundance. We classified the feature table construction methods based on their ability to produce a common feature ID across multiple hypervariable regions (closed reference OTU clustering; n = 13) or not (ASVs and de novo OTUs; n = 19); whether one (n = 10) or multiple (n = 22) hypervariable regions were combined, and whether a specific analysis was done at the OTU/ASV level or higher. In the subset of 32 well defined studies, there was not a statistically significant association with the number of hypervariable regions combined and whether alpha diversity (n=26; Fisher’s p=0.64), beta diversity (n=25; Fisher’s p = 0.39) or differential abundance (n=30; Fisher’s p=0.53) was conducted. There was no significant difference in the feature table construction method based on whether alpha diversity (Fisher’s p=0.36) or differential abundance (Fisher p=1.00) were performed, but beta diversity was conducted in all the studies using closed reference clustering but significantly fewer studies using other techniques (Fisher p=0.02). This was driven primarily by studies using *de novo* clustering, where only half performed beta diversity analysis (Table S1).

We also evaluated whether there was an association between the feature table construction method, number of hypervariable regions, and level at which analyses were conducted. We did not find a statistically significant difference in the relationship between the table construction method and whether analyses were conducted at the feature level for alpha diversity, beta diversity, nor differential abundance. However, we found that the number of hypervariable regions was associated with the taxonomic level used for analysis for beta diversity and differential abundance analyses, but not alpha diversity (Table 5). In both cases, these differences were primarily driven by differences in the number of regions combined when de novo OTUs or ASVs were used. In these cases, feature-based analysis was more common in single region analyses, while taxonomic collapse was more common in multi-region analysis. In contrast, the same was not seen in closed reference OTU clustering, which may reflect the latter’s ability to provide common OTU identifiers across multiple regions, facilitating their comparison.

**Table 5.**
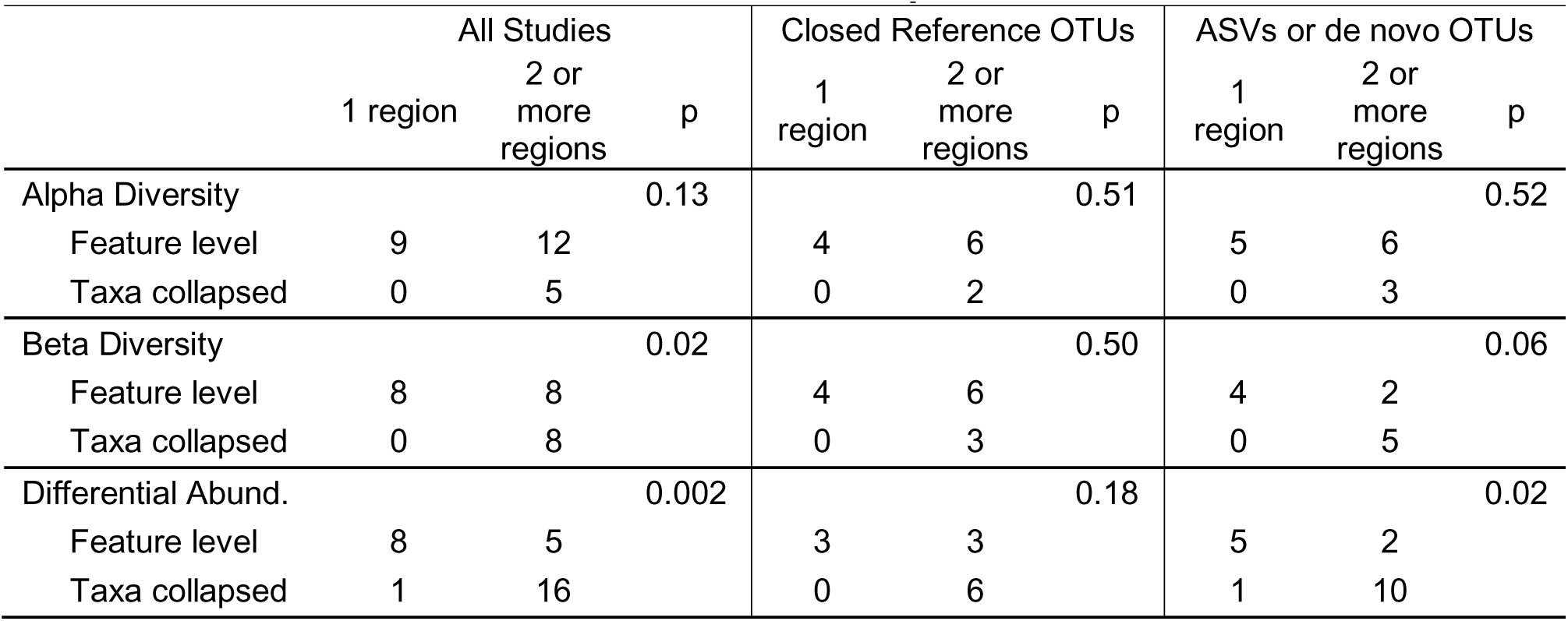
The relationship between the number of hypervariable regions combined, table construction method, and taxonomic collapse.

|  | All Studies |  |  | Closed Reference OTUs |  |  | ASVs or de novo OTUs |  |  |
| --- | --- | --- | --- | --- | --- | --- | --- | --- | --- |
|  | 1 region | 2 or more regions | p | 1 region | 2 or more regions | p | 1 region | 2 or more regions | p |
| Alpha Diversity |  |  | 0.13 |  |  | 0.51 |  |  | 0.52 |
| Feature level | 9 | 12 |  | 4 | 6 |  | 5 | 6 |  |
| Taxa collapsed | 0 | 5 |  | 0 | 2 |  | 0 | 3 |  |
| Beta Diversity |  |  | 0.02 |  |  | 0.50 |  |  | 0.06 |
| Feature level | 8 | 8 |  | 4 | 6 |  | 4 | 2 |  |
| Taxa collapsed | 0 | 8 |  | 0 | 3 |  | 0 | 5 |  |
| Differential Abund. |  |  | 0.002 |  |  | 0.18 |  |  | 0.02 |
| Feature level | 8 | 5 |  | 3 | 3 |  | 5 | 2 |  |
| Taxa collapsed | 1 | 16 |  | 0 | 6 |  | 1 | 10 |  |

## Discussion

We performed a scoping review of English language peer-reviewed studies of the human microbiome sequenced with the 16S rRNA gene that combined multiple data sets. We found that most of the studies were focused on a relationship between the microbiome and an exposure or an outcome, although less than half of these studies used a systematic approach to data set identification, and few described the characteristics of the data sets included. This has the potential to create unobserved bias which can decrease confidence in the conclusion findings of the studies. About a third of studies used open-source data alone, with the remaining two thirds functioning as either consortia or requiring the authors to request data from the original source. This suggests that while there may be a robust open data ecosystem in microbiome science, more work may be needed. Data access may also be a source of bias in the studies.

Finally, we found closed reference OTU clustering was the most common method of combining hypervariable regions, followed by ASV tables, and de novo clustering. While the number of hypervariable regions included was not associated with the feature table construction method, studies using ASVs or performing de novo OTU clustering – methods which were unable to provide common feature-level identifiers across hypervariable regions – were more likely to have performed taxonomically collapsed analyses in beta diversity and differential abundance.

One aim of our work was to summarize the re-use of publicly available data in studies that combined microbiome data sets. In the past decade, there have been increased calls for data sharing both among microbiome scientists (37,38); from funding agencies (48,49); and across scientific disciplines, although adherence is often field-specific (50–52). Proponents of publicly available microbiome data argue that it improves reproducibility, allows innovation, amortizes resources, facilitates better training, and can improve research equity (38,53–55). However, there are concerns about data quality, participant privacy, data sovereignty, “scooping”, and feasibility (55–57). Our scoping review provides insight into data re-use patterns across multiple human microbiome studies, with the caveat that human samples come with additional privacy concerns compared to model organisms or environmental samples. While more than two thirds of studies were able to use some open-source data, only 21 studies used only publicly available data. Nineteen studies had to contact the original authors for additional data. We are unsure whether we should consider it a success that these studies were able to use the restricted access data, since this is often not the case (38,51,58). Furthermore, it is possible that the studies which focused on open data sources also requested data from authors, but were unable to include it due to a lack of response or time restrictions. Both overall and in the context of absent or non-systematic study inclusion criteria, the process of requesting data from authors may also create an unobserved bias, since relevant data sets may have been lost because they could not be accessed. Our results suggest work is needed to explore the impact of using public data only versus requesting data sets from the original authors on both observed associations and the generalizability of findings. Additionally, it suggests that continued support is needed to help achieve the goals of full data sharing to facilitate open collaboration and re-analysis.

One of the surprising themes findings in both our data extraction and conversations within our interdisciplinary team surrounded the use meaning of the word “meta-analysis. There were two major points of disagreement in this discussion: first, whether a meta-analysis required a systematic identification of data sets, and second, whether meta-analysis was restricted to a combination of summary statistics or could be applied to work involving re-analysis of primary data. Many researchers in other fields like genetics, epidemiology, and medicine, have a much stricter concept of “meta-analysis.” The narrowest definitions define a “meta-analysis” as a statistical aggregation of data set-specific estimates (59). Some expanded definitions to allow for other statistical methods, for example, within the individual participant data framework (60,61).

Individual participant data meta-analysis also expands to the approach to allow the analysis of outcomes not presented in the original studies (60,61). Geneticists, in contrast, disambiguate analyses of primary data as “mega-analyses” (62). While not strictly necessary under the restricted statistical framework definition, in many fields the idea of a meta-analysis is intimately tied to a rigorous, systematic approach to data set inclusion, description and analysis in many fields. The Cochrane guidelines (59,60) and others (63,64) caution researchers against conducting meta-analyses without a systematic approach, including a clear search approach, inclusion and exclusion criteria, and strict adherence to a pre-registered analytical protocol (59,60). The systematic, prescribed approach is designed to facilitate a more comprehensive understanding of the review topic; limit bias; enhance transparency; and improve replicability of the review process (65). For many readers, the term “meta-analysis” presupposes this process and allows them to effectively evaluate the thoroughness and comprehensiveness of the evidence included and conclusions drawn. In contrast, we found four prevailing definitions of “meta-analysis” in the microbiome literature. These ran the gamut from a re-analysis of data sets generated by sub studies within a larger parent project to a systematic search, extraction of estimates, and statistical aggregation of results (Table 6). We note that only the last definition encompasses a statistical technique to combine effect estimates across data sets, while the other three approaches may leverage a variety of statistical techniques (or may not conduct statistical testing at all) on sequencing data itself. We acknowledge that each of these designs bring strengths and weaknesses, although we caution extreme care with the fourth design due to strong potential for methodological bias and overwhelming heterogeneity (8–11,18,19). Furthermore, our results suggest that systematic searches are often not performed in microbiome research. Although we did not explicitly analyze the statistical techniques used, re-analysis of primary data is more common than statistically summaries in most cases. We hesitate to recommend terminology, beyond the suggestion that “meta-analysis” be used judiciously; if meta-analysis is expanded beyond the strict statistical definition, microbiome literature might benefit from disambiguation between aggregate statistics and re-analysis of primary data.

**Table 6.** Varying definitions of “meta-analysis” observed in microbiome studies.

|  | Study protocol | Examples |
| --- | --- | --- |
| 1 | Study processes and analyzes data using a common wet lab and bioinformatics protocol, where data were collected for a parent study (e.g., a cohort consortium) that encompasses multiple sub-studies with potentially different designs, study populations, or study sites. | (79,80) |
| 2 | Study combines two or more data sets identified without a systematic search process. Data sets are combined with a common bioinformatic process and analyzed together. Goal of the study is often to provide context or validate findings | (33,35,77,81) |
| 3 | Study identifies data sets through a systematic search process. Data sets are combined through a common bioinformatic process and analyzed together.<br>While not universally true in the current literature, ideally the samples included in the data set are described in a consistent manner, and analytical techniques are appropriate data set heterogeneity (e.g. adjustment for data set specific effects through a fixed or random effect or the statistical pooling of estimates where appropriate for measurements. | (34,82,83) |
| 4<sup>1</sup> | Study identifies data sets through a systematic search process. Summary statistics are extracted from published articles and combined using a statistical meta-analysis framework. | (84–86) |

<sup>1</sup>This definition was observed during screening, but these articles were not included in our final scoping review since we required all included studies combine primary data sets from multiple sources

Microbiome research would also benefit from the development of a microbiome-specific PRISMA guideline, like the STROBE-metagenomics checklist (66), STORMS checklist (67), the PRISMA EcoEvo extension (68), or PRISMA individual patient data extension (69) to help provide a framework for analysis. These solutions will require community buy-in from funders, researchers, editors, and journals. While we, as a small group of researchers, can propose terms, a cross-community working group would be required to finalize guidelines. In the interim, we recommend readers carefully examine the methods to determine if a study presented as a meta-analysis is, in fact, systematic, whether a systematic approach matters for the application in question, and to consider the quality of evidence it may contribute for their work.

Our work has several strengths. First, we had broad inclusion criteria, meaning we included a large variety of studies. We encompassed nearly a decade of work and found studies with a large variety of study designs, purposes, body sites, hypervariable regions, and methodological approaches. Our work is based on a systematic, structured extraction process to limit bias. Extraction was performed by multiple reviewers with regular check-ins and discussion. Finally, we performed a very granular extraction generating 210 unique items, including specificity around table construction strategies, clustering pipeline, and taxonomic levels of analysis used for analysis. However, our work has some limitations. The goal of this analysis was descriptive: we wanted to understand the ways in which microbiome data sets were combined. This did not evaluate the efficacy of the methods for minimizing study effects in data combination.

While a recently published evaluation may be of interest (70), work based on both real data and simulations is needed to evaluate the best way to combine multiple 16S data sets into a single study, both bioinformatically and statistically. We also suffered from a potential for misclassification in our extraction. Microbiome research is a complex, interdisciplinary field with millions of combinatorial possibilities for analysis. Our team found several methods sections to be sparse, missing key clarifying details. For instance, in some cases, we had to rely on the tool name to infer OTU type. In others, details about statistical analyses were non-existent. Finally, analyses were often complex, especially in cases where authors presented a primary analysis with their own data, followed by a contextual analysis with additional data sets. In these cases, the primary analysis might include different analyses than the combined data portion (e.g. alpha diversity, beta diversity conducted on the primary data set; only machine learning on combined data sets). While we tried to ensure consistency through regular extraction comparisons and check-ins, certain complex analyses may have been miscoded. We may have also have limited insight due to our structured extraction form, which may not perfectly fit all included studies. Additionally, our results are necessarily limited due to the scope of our work. We focused on the microbiome characterized using the 16S rRNA genes in samples from humans. Some of the issues we focused on, like the way multiple short read amplicons are combined, are specific to amplicon sequencing and will not generalize to other methods of microbiome characterization, like metagenomics. Our results may also be less relevant to non-human work, where there may be different concerns around issues like data sharing, metadata harmonization, or taxonomic database coverage of key microbes. Finally, our observations are limited by our search terms and extraction. Our structured approach ensured that we collected substantial information but may have inadvertently excluded uncommon protocols.

In conclusion, we conducted a scoping review of combined studies of the human microbiome. Our work suggests that community engagement is needed to determine what a “meta-analysis” means for microbiome studies, and calls for researchers, journals, editors, and readers to consider this closely. We find that while open-source data is widely used, additional efforts could be made to further adoption. Finally, our work re-enforces previous understanding of feature table construction methods to harmonize data sets with disjoint hypervariable regions and primers but begs additional work into their strengths and weaknesses.

## Supporting information

Supplemental File 1. Search Terms

Supplemental File 2. PRISMA checklist

## Acknowledgements

The authors wish to thank Rob Wright at the Welch Medical library for his help in clarifying search terms. Thanks to Luisa Hugerth for her helpful discussion and meticulous copy editing. We also appreciate useful discussion of what a “meta-analysis” means in microbiome research with the attendees of the 2022 Lake Arrowhead Metagenomics meeting for their useful discussion of what “meta-analysis” means in their microbiome research.

Research reported in this publication was supported by the Environmental influences on Child Health Outcomes (ECHO) Program, Office of the Director, National Institutes of Health, under Award Number U24OD023382 (Data Analysis Center). All analyses for scientific publication were performed by the study statisticians, independently of the sponsor. The study sponsor did not review or approve the manuscript for submission to the journal.

## Author Contributions

JWD, AK, NM, JDW, MC, KS, and GP designed the analysis. LR performed the systematic search with input from JWD and AK. YC, MC, KM, JDW, TL, and JWD reviewed articles. YC, YS, ZL, and JWD performed data extraction. JWD curated extracted data. YC, YS, ZL and JWD analyzed the results. ZL and JWD curated analytical code with help from YC and YS. YC, ZL, and JWD wrote the initial draft with help from YS and KM; AK, KM, and JDW provided critical edits. ZL and AK herded all the cats.

All authors reviewed and approved the final manuscript.

## Supplementary Information

Supplementary File 1. Search Terms

Supplementary File 2. PRISMA scoping review checklist

Supplementary Table 1. Relationship between table construction method and whether beta diversity analyses were conducted

