## Supplemental File 1. Search Terms for "In harmony? A scoping review of methods to combine multiple 16S amplicon data sets"

Search Terms (based on consensus)

**PubMed**	~413 results

("Microbiota"[Mesh:NoExp] OR "Gastrointestinal Microbiome"[Mesh] OR "Microbial Consortia"[Mesh] OR "Periphyton"[Mesh] OR "metagenome"[MeSH Terms] OR “metagenomics” [MeSH] OR “dysbiosis”[MeSH] OR microbiome*[tw] OR microbiota*[tw] OR metagen*[tw] OR bacteria*[tw] OR bacterium[tw] OR microorganism*[tw] OR microflora*[tw] OR "intestine flora"[tw] OR "intestinal flora"[tw] OR "gut flora"[tw] OR "microbial consortia*"[tw] OR "microbial consortium*"[tw] OR **"microbial communit*"[tw] OR** periphyton*[tw] OR dysbiosis*[tw] OR dysbioses*[tw] OR disbiosis*[tw] OR disbioses*[tw] OR dysbacterios*[tw] OR disbacterios*[tw])

AND

("Sequence Analysis, DNA"[Mesh] OR "RNA, Ribosomal, 16S"[Mesh] OR "16S"[tw] OR 16SrRNA[tw] OR sequencing[tw] OR "Sequence Analys*"[tw] OR "sequence determination"[tw] OR "community genomics"[tw] OR "population genomics"[tw] OR "environmental genomics"[tw])

AND

("Meta-Analysis"[Publication Type] OR "Meta-Analysis as Topic"[MeSH] OR "meta analys*"[tw] OR "meta-analys*"[tw] OR metanalys*[tw] OR "pooled analy*"[tw] OR "pooled-analy*"[tw] OR "multi-study analy*"[tw] OR "multi study analy*"[tw] OR "horizontal data integration"[tw] OR harmoniz*[tw] OR harmonis*[tw])

**Embase	~441 results**

'microflora'/de OR 'intestine flora'/exp OR 'microbial consortium'/exp OR 'periphyton'/exp OR 'metagenome'/exp OR 'metagenomics'/exp OR 'dysbiosis'/exp OR (microbiome* OR microbiota* OR metagen* OR bacteria* OR microorganism* OR microflora* OR 'intestine flora' OR 'intestinal flora' OR 'gut flora' OR 'microbial consortia*' OR 'microbial consortium*' OR periphyton* OR dysbiosis* OR dysbioses* OR disbiosis* OR disbioses* OR dysbacterios* OR disbacterios*):ab,ti,kw

AND

('DNA sequencing'/exp OR 'RNA 16S'/exp) OR ("16S" OR 16SrRNA OR sequencing OR "Sequence Analys*" OR "sequence determination"):ab,ti,kw

AND

('meta analysis'/exp) OR ("meta analys*" OR "meta-analys*" OR metanalys* OR "pooled analy*" OR "pooled-analy*" OR "multi-study analy*" OR "multi study analy*" OR "horizontal data integration" OR harmoniz*):ab,ti,kw

**Cochrane Library**	(54: 36 reviews, 5 protocols, 13 trials)

https://www.cochranelibrary.com/advanced-search/search-manager?search=6982314

**Scopus**	629 results

TITLE-ABS-KEY ( ( microbiome* OR microbiota* OR metagen* OR bacteria* OR microorganism* OR microflora* OR "intestine flora" OR "intestinal flora" OR "gut flora" OR "microbial consortia*" OR "microbial consortium*" OR periphyton* OR dysbiosis* OR dysbioses* OR disbiosis* OR disbioses* OR dysbacterios* OR disbacterios*) AND ( "16S" OR 16SrRNA OR sequencing OR "Sequence Analys*" OR "sequence determination" ) AND ( "meta analys*" OR "meta-analys*" OR metanalys* OR "pooled analy*" OR "pooled-analy*" OR "multi-study analy*" OR "multi study analy*" OR "horizontal data integration" OR harmoniz* ) )

**Web of Science**	469 results

TS=( ( microbiome* OR microbiota* OR metagen* OR bacteria* OR microorganism* OR microflora* OR "intestine flora" OR "intestinal flora" OR "gut flora" OR "microbial consortia*" OR "microbial consortium*" OR periphyton* OR dysbiosis* OR dysbioses* OR disbiosis* OR disbioses* OR dysbacterios* OR disbacterios*) AND ( "16S" OR 16SrRNA OR sequencing OR "Sequence Analys*" OR "sequence determination" ) AND ( "meta analys*" OR "meta-analys*" OR metanalys* OR "pooled analy*" OR "pooled-analy*" OR "multi-study analy*" OR "multi study analy*" OR "horizontal data integration" OR harmoniz* ) )
