## Supplemental Table 1 for "In harmony? A scoping review of methods to combine multiple 16S amplicon data sets"

Table S1. Relationship between table construction method and whether beta diversity analyses were conducted

|  | Closed ref OTUs | ASVs | De novo OTUs | ASVs or de novo OTUs |
| --- | --- | --- | --- | --- |
| Beta Diversity |  |  |  |  |
| No | 0 | 4 | 3 | 7 |
| Yes | 13 | 9 | 3 | 12 |
